# The Secretome of Liver X Receptor Agonist Treated Early Outgrowth Cells Decreases Atherosclerosis in *Ldlr*-/- Mice

**DOI:** 10.1101/757435

**Authors:** Adil Rasheed, Sarah Shawky, Ricky Tsai, Richard G Jung, Trevor Simard, Benjamin Hibbert, Katey J Rayner, Carolyn L Cummins

**Author notes:** Please address correspondence to: Carolyn L. Cummins, Faculty of Pharmacy, University of Toronto, 144 College Street Room 1101, Toronto, Ontario, M5S 3M2.

## Abstract

**Objective:** Endothelial progenitor cells (EPCs) promote the maintenance of the endothelium by the secretion of vasoreparative factors. A population of EPCs known as early outgrowth cells (EOCs) are currently being investigated as novel cell-based therapies for the treatment of cardiovascular disease. We previously demonstrated that the absence of liver x receptors (LXRs) is detrimental to the formation and function of EOCs under hypercholesterolemic conditions. Here, we investigate whether LXR gain-of-function in EOCs is beneficial for the treatment of atherosclerosis.

**Approach and Results:** EOCs were differentiated from the bone marrow of wildtype (WT) and LXR-knockout (*Lxrαβ*-/-) mice in the presence of vehicle or LXR agonist (GW3965). WT EOCs treated with GW3965 throughout differentiation showed reduced expression of endothelial lineage markers (*Cd144, Vegfr2*) compared to WT vehicle and *Lxrαβ*-/- cells. GW3965-treated EOCs produced secreted factors that reduced monocyte adhesion to activated endothelial cells in culture. When injected into atherosclerosis-prone *Ldlr*-/- mice, GW3965-treated EOCs and concentrated conditioned media (CM) from GW3965-treated EOCs, reduced plaque burden within the aortic sinus. Furthermore, when CM from human EOCs (obtained from patients with established CAD) were treated with GW3965, monocyte to endothelial adhesion was decreased suggesting the translatability of the results.

**Conclusions:** *Ex vivo* LXR agonist treatment of EOCs produces a secretome that decreases early atherosclerosis in *Ldlr*-/- mice. CM from human EOCs significantly inhibits monocyte to endothelial adhesion. Thus, active factor(s) within the GW3965-treated EOC secretome have the potential to be useful for the treatment of atherosclerosis.

## Introduction

Cardiovascular disease remains the leading cause of death worldwide.^1^ Atherosclerosis is a vascular complication arising from cardiovascular disease and is typically diagnosed in its later stages, when established lipid-laden plaques are deposited in the aorta, narrowing the vessel wall and resulting in ischemia. Plaque development progresses through multiple stages over many decades.^2^ However, development of these plaques is initiated by coordinate dysfunction to various vessel wall cell types, including endothelial and vascular smooth muscle cells, as well as hematopoietic cell types.^2^ Prior to plaque deposition, the aortic endothelium undergoes pathological activation resulting from systemic inflammation and elevations in plasma lipoproteins. During endothelial activation, selectins and adhesion molecules are upregulated, which facilitate the binding of monocytes to the endothelium. Adherent monocytes then transverse the endothelium to the intima where they differentiate into macrophages and then engulf modified lipoproteins. These lipid-laden foam cells promote the development of plaques which are characteristic of established atherosclerosis.^3^

The liver x receptors, LXRα (*Nr1h3*) and LXRβ (*Nr1h2*), belong to the nuclear receptor superfamily of transcription factors.^4, 5^ Activation of the LXRs induces the gene expression of the cholesterol efflux transporters, *Abca1* and *Abcg1*; and represses pro-inflammatory gene expression, including *Mcp-1, Tnfα*, and *Il-1β*.^6-8^ These roles of LXRs have been particularly well characterized in macrophages, in part through bone marrow transplant experiments, that support an anti-atherogenic role for LXRs.^9-14^ However, recent bone marrow transplantation studies using bone marrow deficient in LXR target genes have demonstrated that LXRs may elicit their anti-atherogenic roles in other bone marrow-derived cell types apart from monocytes/macrophages.^14-17^ In the bone marrow, hematopoietic stem cells (HSCs) can differentiate into myeloid progenitors, which produce the monocytes/macrophages known to contribute directly to plaque progression.^3, 18^ HSCs, however, can also differentiate into other cell types including endothelial progenitor cells (EPCs).^19, 20^

EPCs were initially described in a report by Asahara *et al.* in 1997 as peripheral blood cells that differentiate into endothelial-like cells and contribute to vascular repair by direct incorporation.^19^ Patients with diabetes and cardiovascular disease have demonstrated dysfunction in their EPC numbers and function.^21-23^ Over the past 20 years, EPC function has largely been characterized using *ex vivo* cultures, which yield two cell populations: early outgrowth cells (EOCs; 7 days in culture) and late outgrowth endothelial cells (21 days in culture). Consistent with observations made by Asahara and colleagues, many believe that EPCs exert their function by acquiring an endothelial-like phenotype and facilitate neovascularization by direct incorporation.^24-29^ However, studies have shown inconsistent engraftment *in vivo*, suggesting that endothelial repair may occur via an indirect mechanism.^30-34^ Given these discrepancies, attention has shifted to a mechanism of action whereby EOCs support endothelial repair indirectly by the secretion of vasoreparative factors. These EOCs have been of particular interest in recent years spurring a new generation of clinical trials^35, 36^ and applications in regenerative medicine reviewed by Chong *et al*. (2016).^37^

Using *Lxrαβ*-/-double knockout mice fed a hypercholesterolemic diet, we previously demonstrated that LXRs are essential for preventing cholesterol-induced defects in EOCs.^38^ Elevated cholesterol content only in the absence of LXRs, altered EOC differentiation and the resulting secretome, which increased monocyte adhesion to treated endothelial cells *in vitro*. As a result of these studies, we set out to determine: i) whether pharmacologic activation of LXRs could beneficially influence EOC differentiation and secretome function, and ii) if *ex vivo* pharmacological activation of LXRs in EOCs could prevent the development of atherosclerosis *in vivo*. Herein, we present data that show that pharmacological activation of LXRs reduces the expression of endothelial lineage markers in WT EOCs, and alters the EOC secretome in a manner that inhibits monocyte adhesion to treated endothelial cells. Tail vein administration of either LXR agonist-treated WT EOCs or their concentrated conditioned media to *Ldlr*-/- mice attenuated lesion development in the aortic sinus compared to their respective controls. Furthermore, we demonstrate the translational potential of LXR agonist as a therapeutic using EOCs derived from patients with established coronary artery disease (CAD). The studies presented here suggest that activation of LXRs in EOCs *ex vivo* may provide a new avenue for therapeutic intervention in atherosclerosis.

## Materials and methods

Materials and Methods are available in the online-only Data Supplement.

## Results

### GW3965 alters differentiation of EOCs to the endothelial lineage

We previously demonstrated using a loss-of-function model that LXRs play an important role in mitigating the negative effects of cholesterol, either from Western diet feeding or direct addition of exogenous cholesterol, on EOC differentiation.^38^ Here we set out to determine whether LXR activation would have an opposing role on EOC differentiation. Treatment with LXR agonist (1 μM GW3965) throughout EOC differentiation decreased the expression of the endothelial lineage markers *Cd144* and *Vegfr2* both by 76% in day 7 EOCs derived from WT mice, but not *Lxrαβ*-/- mice (**Figure 1**). Interestingly, GW3965 treatment resulted in a varied time course profile for these endothelial genes. WT cells differentiated in the presence of GW3965 showed similar expression of *Cd144* compared to vehicle treated cells up to day 3. However, after day 3, *Cd144* expression was decreased in the WT GW3965-treated EOCs compared to vehicle treatment, suggesting a de-differentiation program (**Figure 1A**). On the other hand, the increase in *Vegfr2* expression normally observed with differentiation was prevented in WT cells treated with GW3965 (**Figure 1B**).

**Figure 1.**
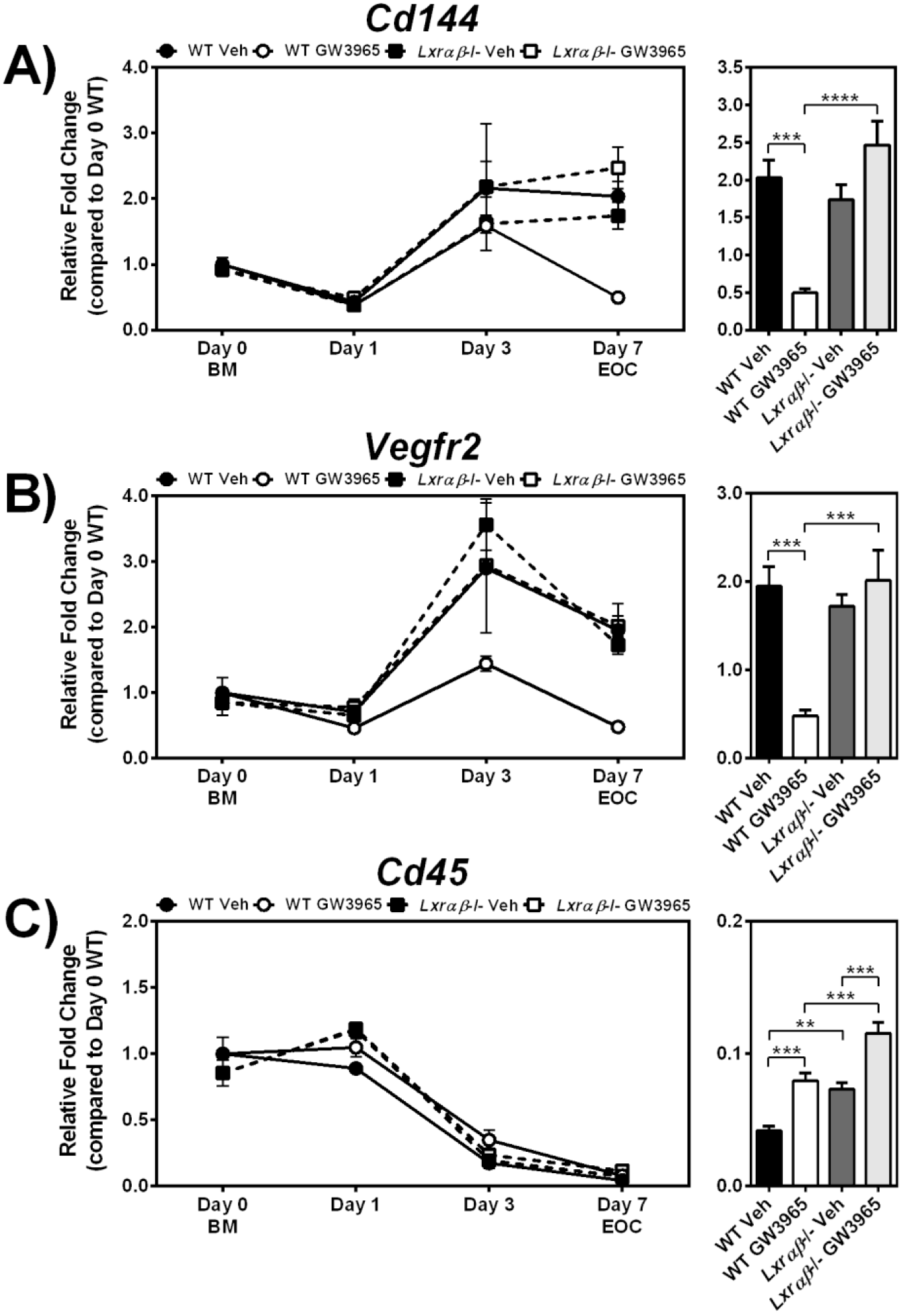
Wildtype EOCs treated with LXR agonist have lower expression of endothelial markers after differentiation. Bone marrow (BM; Day 0) from 12-week old WT and *Lxrαβ*-/- mice were differentiated for 7 days in the presence of LXR agonist (1 μM GW3965). Gene expression was performed for the endothelial lineage markers **A**, *Cd144* and **B**, *Vegfr2*, as well as **C**, the leukocyte marker *Cd45*. Time course analyses are shown in the left panels, and day 7 EOC expression levels in the right panel. n=5-6 per group. Data represent the mean ± SEM. \*\**P*<0.01, \*\*\**P*<0.001, \*\*\*\**P*<0.0001.

To determine whether LXR activation in WT EOCs alters the expression of other hematopoietic lineage markers, we next examined the expression of leukocyte/myeloid markers, which are expected to decrease during differentiation. Time course analysis did reveal potent reductions of the pan leukocyte marker *Cd45* by day 3 of differentiation, and while GW3965 treatment appeared to delay this decrease at day 7 in EOCs, this effect was not LXR-dependent (**Figure 1C**). Furthermore, at day 7, expression of the myeloid marker *Cd11b* was not altered by GW3965 in WT EOCs (**Supplemental Figure 1A**). During development, endothelial to mesenchymal transition (EndMT) is critical to ensure proper structural development of the heart, where cardiac endothelial cells give rise to mesenchymal precursors eventually forming valves. However, under pathological states, EndMT induced by oxidative stress, TGFβ, and other characteristic features of the atherosclerotic milieu, can cause endothelial cells to acquire a more fibrotic phenotype. Cells undergoing this transition show increases in mesenchymal markers such as *Tgfβ* and *Fsp-1*.^39, 40^ Notably, while LXR activation decreased EOC expression of endothelial lineage markers (*Cd144* and *Vegfr2*), expression of these mesenchymal markers in the GW3965-treated WT EOCs was not increased (**Supplemental Figure 1B-C**), suggesting that these EOCs were not undergoing EndMT. These data indicate that while GW3965 treatment to WT cells decreased differentiation to the endothelial lineage, expression of the leukocyte/myeloid markers *Cd45* and *Cd11b* were not increased, suggesting that agonist treatment did not prevent differentiation, but rather produced an altered population of EOCs.

### Secretome of GW3965-treated EOCs reduced monocyte endothelial adhesion

To investigate the impact of LXR-activated EOCs during the early stages of atherosclerosis, we used the *in vitro* monocyte-endothelium adhesion assay. Human umbilical vein endothelial cells (HUVECs) activated with TNFα were incubated with conditioned media (CM) from vehicle- or GW3965-treated EOCs prior to the addition of fluorescently labeled THP-1 monocytes. As expected, TNFα-treatment increased monocyte adhesion to HUVECs by 8.8-fold (*P*<0.0001, **Figure 2A**). Pre-treatment of HUVECs with CM from GW3965-treated WT EOCs (WT GW CM) decreased the adhesion of THP-1 cells by 36% compared to WT Veh CM (*P*<0.0001, **Figure 2A**). In contrast, CM from GW3965-treated *Lxrαβ*-/- EOCs (*Lxr*-/- GW CM) had no effect on TNFα-induced monocyte adherence. These results indicate that LXR activation altered the EOC population, which in turn produced a secretome that decreased monocyte adhesion to activated endothelial cells.

**Figure 2.**
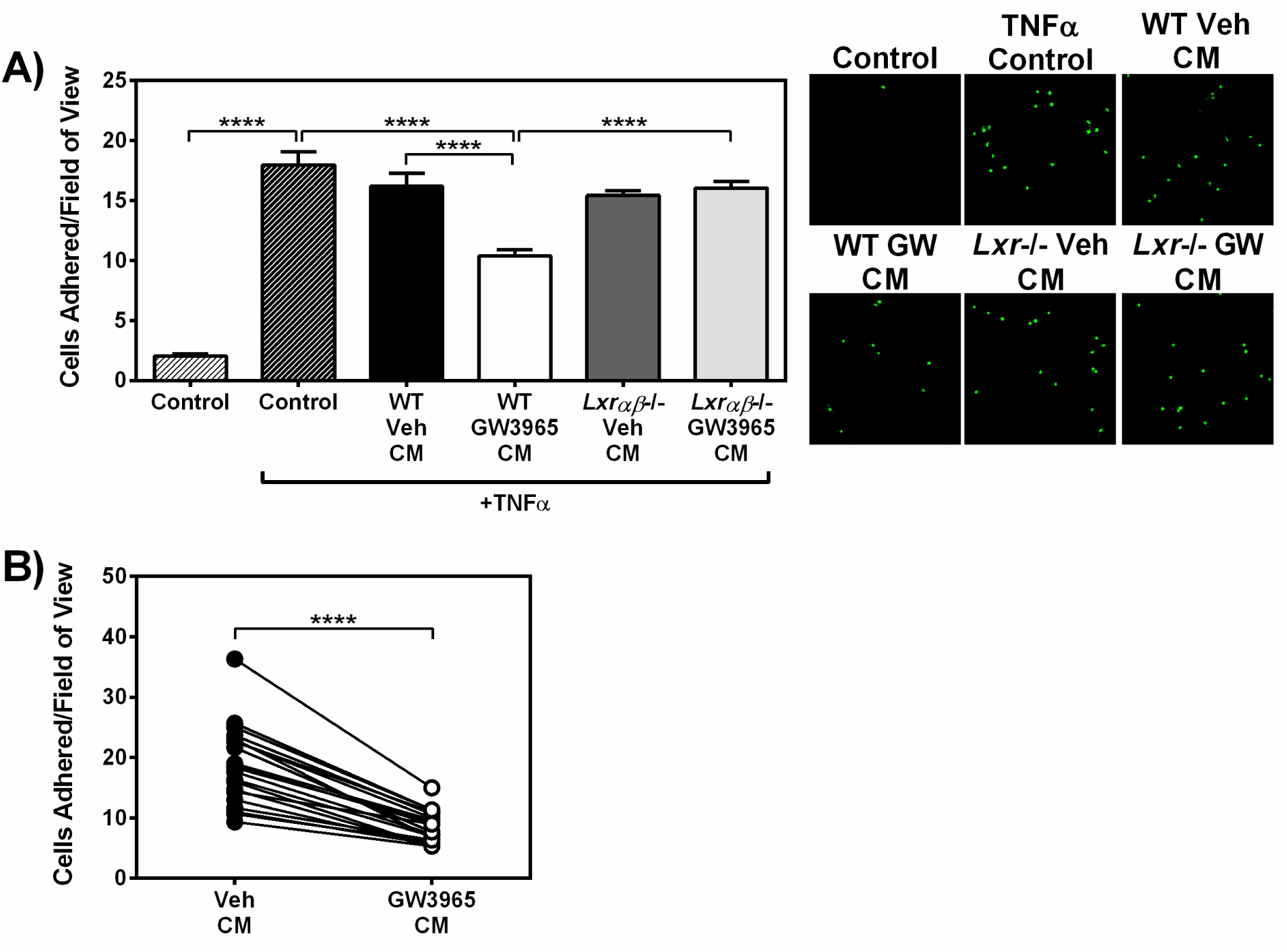
Secreted factors from GW3965-treated mouse and patient-derived EOCs reduce monocyte adherence to activated endothelial cells. Conditioned media from GW3965-treated EOCs were harvested and applied to HUVECs treated with TNFα. Monocyte adhesion to treated HUVECs was quantified after incubation with conditioned media derived from **A**, WT and *Lxrαβ*-/- mouse EOCs (n=8 per group) and **B**, EOCs derived from patients with coronary artery disease (n=21 patients). Data represent the mean ± SEM. \*\*\*\**P*<0.0001.

During atherosclerosis, endothelial cells upregulate the expression of selectins, which facilitate leukocyte rolling, and adhesion molecules that enhance leukocyte adhesion to the aortic endothelium.^3^ To determine whether the WT GW CM was decreasing monocyte adhesion by altering the expression of adhesion molecules via signaling to the HUVECs, we performed gene expression analysis of the HUVECs after exposure to EOC CM. We observed that while the addition of the pro-inflammatory stimulus TNFα did induce the expression of the adhesion molecules *VCAM1* and *ICAM1*, as well as *SELE*, there was no change in any of these markers of endothelial activation when co-treated with WT GW CM (**Supplemental Figure 2**). As such the mechanism by which WT GW CM alters monocyte-HUVEC adhesion remains elusive.

We set out to determine whether the therapeutic effects of the GW3965 on the EOC secretome could be translated to patients with coronary artery disease (CAD). For this, we obtained peripheral blood samples from a CAD cohort with characteristics described in **Table 1**. Cells were differentiated to EOCs in the presence of vehicle or 1 μM GW3965 and the CM was collected at day-7. Similar to EOCs derived from mouse bone marrow, monocyte adhesion was significantly decreased by 54% when HUVECs were treated with the CM from GW3965-treated EOCs from the CAD patients compared to vehicle treatment (**Figure 2B**). Our data indicate that GW3965 improved EOC function from CAD patients to a similar extent as that observed in murine cells from WT mice.

**Table 1.** Characteristics of coronary artery disease patients from which EOCs were derived.

|  | EOC (n = 21) |  |
| --- | --- | --- |
| | Number or mean ± SD | Proportion (%) |
| **Age** | 68 ± 13 | |
| **Sex (male)** | 15 | 71.4 |
| **HTN** | 12 | 57.1 |
| **DM** |  |  |
| *Type I* | 1 | 4.8 |
| *Type II - oral agents* | 4 | 19.1 |
| *Type II - insulin* | 2 | 9.5 |
| **Smoker** |  |  |
| *Never* | 16 | 76.2 |
| *Remote (quit >1 month ago)* | 3 | 14.3 |
| *Active* | 2 | 9.5 |
| **Dyslipidemia** | 12 | 57.1 |
| **Family history of CAD** | 3 | 14.3 |
| **ACS** | 7 | 33.3 |
| **HF/LV dysfunction** | 2 | 9.5 |
| **Prior History** |  |  |
| *MI* | 4 | 19.1 |
| *PCI* | 3 | 14.3 |
| *CVA* | 1 | 4.8 |
| *CABG* | 1 | 4.8 |
| **Medications** |  |  |
| *ASA* | 20 | 95.2 |
| *P2Y12* | 18 | 85.7 |
| *Beta blocker* | 10 | 47.6 |
| *ACEi/ARB* | 14 | 66.7 |
| *Statin* | 19 | 90.5 |
| **Biochemistry** |  |  |
| *WBC (10<sup>9</sup> cells/L)* | 7 ± 2 | |
| *HGB (g/L)* | 134 ± 26 | |
| *PLT (10<sup>9</sup> cells/L)* | 225 ± 52 | |
| *HbA1c (%) (n = 16)* | 6.1 ± 0.8 | |
| *Cholesterol (mM) (n = 7)* | 4 ± 1 | |
| *LDL (mM) (n = 7)* | 2 ± 1 | |
| *Creatinine (μM)* | 92 ± 65 | |

ACE: angiotensin-converting enzyme; ACS: acute coronary syndrome; ARB: angiotensin receptor blocker; ASA: acetylsalicylic acid; CABG: coronary artery bypass graft; CVA: cerebrovascular accident; DM: diabetes mellitus; HF/LV: heart failure/left ventricular; HGB: hemoglobin; HTN: hypertension; LDL: low-density lipoprotein; MI: myocardial infarction; PCI: percutaneous coronary intervention; PLT: platelet; WBC: white blood cell

### Administration of EOCs and their secreted factors reduce plaque size during atherogenesis

*Ex vivo* cultured EOCs are currently undergoing clinical trials for the treatment of vascular complications arising from cardiovascular disease, such as pulmonary hypertension and acute myocardial infarction.^35-37^ In our study, we wanted to assess the therapeutic potential of WT GW3965-treated EOCs and their secretome specifically with respect to the development of atherosclerosis. We used atherosclerosis-prone *Ldlr*-/- mice randomized to receive WT EOCs differentiated in the presence of either Veh or GW3965 and a separate cohort that was randomized to receive only the concentrated CM from these cell treatments. The 8-week-old *Ldlr*-/- mice from either cohort were fed a Western diet for 8 weeks, during which the EOCs or CM was administered every 2 weeks.

The *Ldlr*-/- mice receiving EOCs were splenectomized prior to initiation of the treatments to prevent sequestration of EOCs within the spleen.^41^ Freshly cultured EOCs were administered to the *Ldlr*-/- mice bi-weekly throughout the Western diet feeding period (**Figure 3A**). EOC administration did not alter plasma cholesterol or circulating inflammatory cell types of *Ldlr*-/- mice compared to saline injected mice (**Figure 3B & Supplemental Table 3**). Histological analysis of the aortic sinus revealed a reduction in the luminal plaque coverage of the *Ldlr*-/- mice receiving the GW3965-treated EOCs compared to those receiving the Veh-treated EOCs (44%, *P*<0.05) or saline (41%, n.s.) (**Figure 3C-D**). Since our *in vitro* results found a potent effect of the EOC secretome on limiting monocyte-endothelial adhesion (**Figure 2A**), we administered CM from treated EOCs or control (unconditioned) media bi-weekly to *Ldlr*-/- mice (with an intact spleen) to determine whether this alone would be sufficient to impact lesion formation (**Figure 4A**). We did not observe changes in plasma cholesterol upon injection of the CM (**Figure 4B**). Likewise, there was no effect on total blood counts of mice treated with CM, with the exception of mice receiving CM from GW3965-treated EOCs in which monocytes were unexpectedly increased (**Supplemental Table 4**). Remarkably, treatment with CM from GW3965-treated EOCs reduced atherosclerotic plaques in the aortic sinus by 36% compared to Veh-treated EOC CM and 47% compared to control unconditioned EBM-2 media (**Figure 4C-D**). While the mechanism by which these changes occur is not clear, these data demonstrate the therapeutic potential of the GW3965-treated EOC secretome to limit the development of atherosclerosis in mice, with *in vitro* evidence that this response will be conserved in humans.

**Figure 3.**
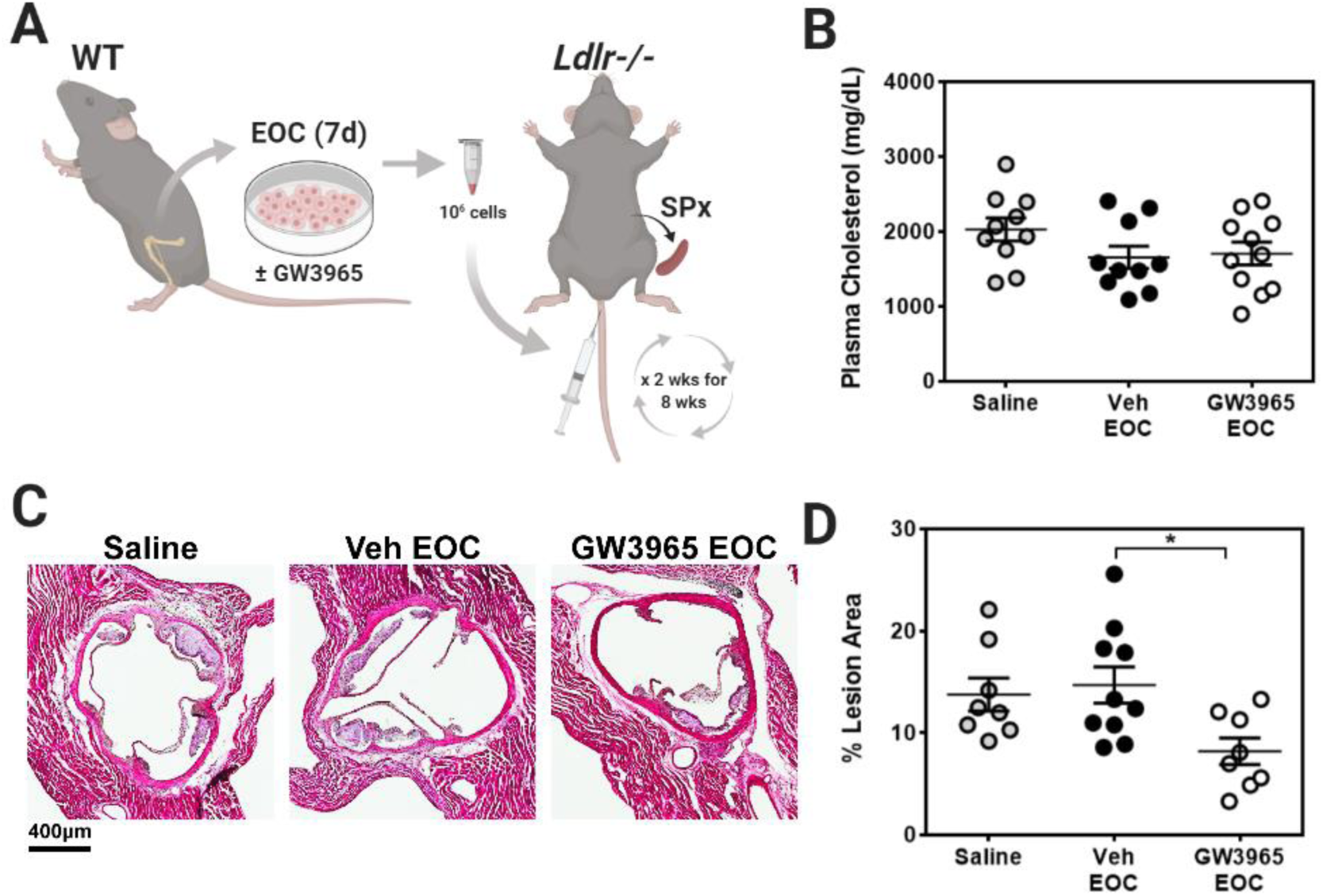
LXR agonist-treated EOCs reduce plaque area in the aortic sinus. **A**, EOCs derived from WT mice were differentiated in the presence of GW3965 and administered to splenectomized *Ldlr*-/- mice bi-weekly over an 8-week period. **B**, Circulating plasma cholesterol levels were quantified from the blood collected at sacrifice. **C**, Aortic sinus sections were stained with H&E and **D**, plaque area was quantified by an experimenter blinded to the treatment condition. n=8-11 mice per group. Data represent the mean ± SEM. \**P*<0.05.

**Figure 4.**
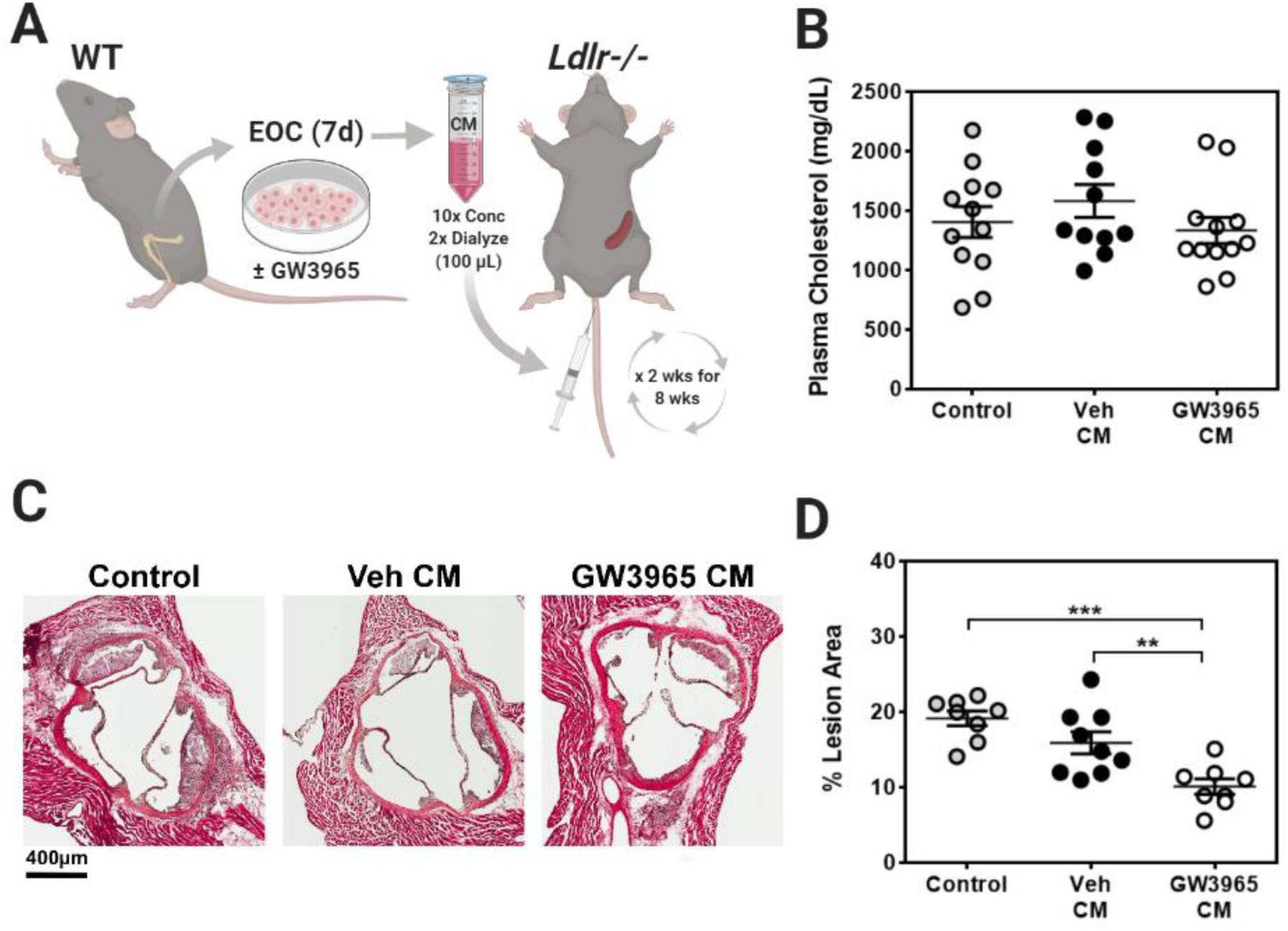
The secretome of GW3965-treated EOCs decreases atherosclerotic plaque burden. **A**, The conditioned media from GW3965 treated EOCs were harvested and administered to *Ldlr*-/- mice bi-weekly over 8 weeks. **B**, Plasma cholesterol levels were assessed in samples collected at sacrifice. **C**, Aortic sinus sections were stained with H&E and **D**, plaque area was quantified by an experimenter blinded to the treatment condition. n=8-12 mice per group. Data represent the mean ± SEM. \*\**P*<0.01, \*\*\**P*<0.001.

## Discussion

Atherosclerosis is a progressive disease that manifests over several decades. The gold standard treatment for atherosclerosis relies on lipid-lowering, via administration of statin drugs and more recently PCSK9 inhibitors.^42^ However, results from the Canakinumab Anti-inflammatory Thrombosis Outcome Study (CANTOS) trial found that targeting the inflammatory axis during cardiovascular disease can reduce cardiovascular-related mortality, independent of lipid lowering.^42, 43^ These data, among others, emphasize the opportunity for the development of novel therapeutic avenues for the treatment of atherosclerosis that go beyond traditional lipid lowering modalities.

Atherosclerosis begins with endothelial defects that, if not repaired, promote the recruitment of immune cells that contribute to the formation of the atherosclerotic plaque. During these early stages, damage that occurs from the atherosclerotic milieu overwhelms endogenous endothelial repair mechanisms, resulting in an overall pro-inflammatory response of the endothelium allowing for an increase in expression of adhesion molecules that facilitate leukocyte binding.^3^ While the endothelium has the capability to repair itself, endothelial repair can also be supported by EOCs, through the secretion of vasoreparative factors. We previously described a role for LXRs in mitigating the negative effects of high cholesterol-induced defects on EOC differentiation and the secretome.^38^ Given the importance of EOCs in mediating endothelial repair via the secretome and the contributions of endothelial activation to the early pathogenesis of atherosclerosis, we tested whether pharmacologic activation of LXRs in EOCs produces a secretome that could reduce the progression of early-stage atherosclerosis. Our data show that LXR activation during EOC differentiation decreases the expression of endothelial lineage markers. This shift in differentiation qualitatively alters the secretome of the GW3965-treated EOCs in a favorable manner, resulting in less monocyte-endothelial adherence and an overall decrease in atherosclerosis. Along with our previously published data^38^, this therefore provides a model whereby expression of endothelial markers is inversely associated with the benefit of the EOC secretome.

In contrast to the loss-of-function model, in which increased cholesterol in the *Lxrαβ*-/- EOCs correlated with increased expression of endothelial lineage markers^38^; the gain-of-function LXR model studied here demonstrated a decrease in these endothelial markers in the absence of changes in EOC cholesterol content (data not shown). These data suggest that there are separate mechanisms regulating EOC differentiation under the influence of cholesterol when LXRs are absent compared to when they are activated. Pharmacologic activation of LXRs throughout differentiation specifically decreased the expression of the endothelial lineage markers *Cd144* and *Vegfr2*. Time course analysis revealed that GW3965 treatment prevented the increase of *Vegfr2*, while *Cd144* expression decreased after day 3 of differentiation, compared to vehicle-treated cells. Further experiments will be needed to confirm the mechanism by which EOC differentiation to the endothelial lineage is impacted by the activation of LXR, and how this is occurring for each endothelial marker. These experiments will need to determine whether LXRs are directly binding to the DNA to inhibit endothelial marker expression or interact with other factors to inhibit transcriptional machinery at these individual loci.

We attempted to narrow down potential mechanisms by which GW3965-treated EOC derived conditioned media decreased monocyte to endothelial adhesion using gene expression analysis of HUVECs. We found no significant differences in marker genes associated with enhanced adhesion; thus, the mechanism by which the secretome exerts this effect is still elusive. Signals transduced from the secretome are likely derived from proteins^44-50^ or components of extracellular vesicles (i.e., miRNAs).^51-53^ As such, further investigation of the LXR-dependent changes to the EOC secretome is warranted to identify potential effector pathways of endothelial cells that lead to decreased monocyte adhesion, including unbiased proteomic approaches and analysis of RNA species contained within the ligand-treated EOC secretome. Nevertheless, we show here that GW3965-treatment of EOCs derived from patients with CAD produces a secretome that also reduces monocyte-endothelial adhesion (**Figure 2B**), indicating that LXR agonist treatment to EOCs could potentially be a translatable therapeutic for patients with cardiovascular disease.

Previous studies have demonstrated that administration of EOCs to atherosclerotic mouse models can promote plaque stabilization and decrease plaque formation, primarily through endothelial engraftment.^54-57^ However, those studies were all performed in mouse models of plaque regression that targeted the later stages of atherosclerosis. EOC intervention at the early stages of atherosclerosis, as described herein, has not, to our knowledge, been previously explored. As a complement to our *in vitro* monocyte-endothelial adhesion assay, we evaluated the therapeutic potential of LXR treatment to EOCs in a mouse model of early atherosclerosis. In line with numerous studies and clinical trials (reviewed by Mindur and Swirsky (2019)^58^) that show the potency of circulating growth factors and cytokines in promoting and/or limiting the development of atherosclerosis, administration of the secreted factors from GW3965-treated EOCs significantly reduced aortic lesion development in recipient *Ldlr*-/- mice. Of note, the effect of injecting GW3965-treated EOCs vs. GW3965-treated CM on lesion formation was very similar (44% vs. 36%, respectively) indicating that the majority of the beneficial effect is derived from the secretome. These data are in line with recent reports in which the EOC secretome has been packaged in nanoparticles or hydrogels to treat acute ischemic diseases.^59, 60^

Previous studies determined that in the absence of the cholesterol efflux transporters (*Abca1* and *Abcg1*) in myeloid cells of the bone marrow was insufficient to explain the anti-atherogenic effects of LXRs described in whole bone marrow transplant experiments.^9-14^ Our data support the role of another effector cell (EPC) in contributing to the anti-atherogenic effects of LXR. Thus, these studies enhance our understanding of LXR cell targets in the bone marrow and provide a unique mechanism to inhibit the development of atherosclerosis. A major drawback to the *in vivo* use of LXR agonists is the development of hypertriglyceridemia and hepatosteatosis via upregulation of the LXR target gene and lipogenic factor *Srebp1c*, in the liver.^61-63^ Our data demonstrate that the cell-free secretome of LXR-treated EOCs is sufficient to protect against endothelial damage and early atherosclerosis. These data therefore support a role for *ex vivo* treatment of patient-derived EOCs as an avenue of intervention that would elicit the anti-atherogenic effects of LXRs in the EOCs without the hepatosteatosis associated with systemic LXR agonism.

## Acknowledgments

We would like to sincerely thank Dr. David J. Mangelsdorf (University of Texas Southwestern Medical Center, Dallas, TX) for kindly providing the Liver X Receptor knockout mice, Dr. Erin E. Mulvihill for her guidance with the animal sacrifices, and Xiaoling Zhao for processing samples for histology.

## Highlights

- LXR agonist treatment during EOC differentiation reduces endothelial marker expression and produces a secretome that represses monocyte adhesion to activated endothelial cells in culture.
- LXR agonist-treated EOCs derived from CAD patients also secrete factors that reduce monocyte-endothelial adhesion.
- Administration of the conditioned media from LXR agonist-treated EOCs significantly reduces plaque development in recipient *Ldlr*-knockout mice.

## Abbreviations

CAD: coronary artery disease
CM: conditioned media
EndMT: endothelial-to-mesenchymal transition
EOC: early outgrowth cell
EPC: endothelial progenitor cell
HUVEC: human umbilical vein endothelial cell
LXR: liver x receptor
WT: wildtype

## SUPPLEMENTAL MATERIAL

### Materials and Methods

#### Mice

All animal procedures were approved by the Institutional Animal Care and Use Committee at the University of Toronto. WT and LXR double knockout (*Lxrαβ*-/-) male mice were backcrossed more than 10 generations on a C57Bl/6 background and maintained on a chow diet (#2016 Harlan Teklad, Mississauga, ON, Canada). *Ldlr*-knockout male mice (*Ldlr*-/-; B6.129S7-Ldlr^*tm1Her*^/J, Stock #002207) on a C57Bl6/J background were purchased from Jackson laboratories (Bar Harbor, ME). All mice were housed in a temperature and light-controlled environment. Mice were sacrificed between 9am and 11am by cervical dislocation under isoflurane anesthesia.

#### Bone marrow harvesting and early outgrowth cell culture

EOCs were cultured from the bone marrow of the tibiae and femurs of WT and *Lxrαβ*-/- mice. The bone marrow cells were seeded on human-fibronectin coated (Sigma-Aldrich, Oakville, ON, Canada) tissue culture dishes and differentiated for 7 days to EOCs in endothelial basal media supplemented with growth factors/cytokines (EGM-2 Bullet Kit; Lonza, Walkersville, MD) at 37°C with 5% CO_2_. The media was changed every other day. EOCs were cultured in the presence of vehicle (ethanol) or LXR agonist (1 μM GW3965; Tocris, Minneapolis, MN) throughout the 7-day differentiation protocol.

#### RNA isolation, cDNA synthesis, and real-time QPCR

RNA isolation, cDNA synthesis, and QPCR were performed as previously described^38^. Primer sequences are listed in Supplemental Table 1 (mouse) and Supplemental Table 2 (human).

#### Coronary artery disease patient recruitment

Patients who underwent coronary angiography gave written informed consent for blood collection. This study was approved by the Ottawa Health Science Network Research Ethics Board (OHSN-REB, Protocol #: 20160516-01H). This study conforms with the 1975 Declaration of Helsinki for the use of human blood. The University of Ottawa Heart Institute serves 1.2 million people with all cardiac catherization procedures prospectively indexed in the CArdiovascular and Percutaneous clInical Trials (CAPITAL) revascularization registry along with baseline investigations and medications.^64, 65^ From August 2018 to March 2019, blood samples were collected from 21 patients in EDTA tubes (Becton Dickinson, Franklin Lakes, NJ) at the time of coronary angiography. Coronary artery disease was confirmed by coronary angiography and defined as an epicardial stenosis ≥50%. The presence of type 2 diabetes was defined as hemoglobin A1c (HbA1c) levels ≥6.5% or a prior diagnosis or medical therapy for diabetes. Patient characteristics are provided in Table 1.

#### Human EOC culture

Peripheral blood mononuclear cells were isolated by density centrifugation using Ficoll-Paque Plus (GE Healthcare, Marlborough, MA). The cells were differentiated to EOCs as described above. For each patient, the mononuclear cells were differentiated to EOCs in both vehicle and 1 μM GW3965.

#### Conditioned media collection

After 7 days of differentiation, EOCs were washed twice with cold PBS and incubated in factor- and GW3965-free media (EBM-2; Lonza) for 30 minutes at 37°C and 5% CO_2_. The media was discarded and the cells were replenished with fresh EBM-2 media. The conditioned media (CM) was then collected after 24 hours, centrifuged at 700*xg* for 5 minutes to remove any cells, and syringe filtered (0.22 μm). The CM was frozen at -80°C until use. The number of EOCs from each treatment group was unchanged after differentiation (data not shown).

#### Cell lines

Human umbilical vein endothelial cells (HUVEC) and media (EGM-2 Bullet Kit) were purchased from Lonza. THP-1 monocytes were received as a gift from Dr. Myron I. Cybulsky (University of Toronto) and were maintained in RPMI 1640 with _L_-glutamine (Sigma-Aldrich) supplemented with 10% FBS and 0.05 mM β-mercaptoethanol (Sigma-Aldrich). All cell lines were used between passages 4-6.

#### Monocyte-endothelial adhesion assay

HUVECs were seeded in 12-well plates on sterilized circular glass coverslips (Fisher Scientific, Hampton, NH) and treated at 60% confluence with 20% (mouse) or 30% (human) EOC CM for 20 hours in EGM-2 media, followed by a 4-hour co-treatment with EOC CM and 10 ng/mL TNFα (Life Technologies). After treatment, HUVECs were co-incubated with 10^5^ CMFDA (Life Technologies) -labelled THP-1 monocytes per well for 90 minutes. After co-culture, the THP-1 cells were washed off and adherent THP-1 cells were fixed using 4% paraformaldehyde. Coverslips were mounted onto microscope slides using Vectashield (Vector Labs, Burlingame, CA) or Dako Fluorescence (Agilent Technologies, Santa Clara, CA) and images were acquired on a laser confocal LSM700 or LSM880 microscope operated by Zen software (Zeiss, Toronto, ON, Canada).

#### Splenectomy

A cohort of *Ldlr*-/- mice were splenectomized at 4 weeks of age. Under isoflurane anesthesia, the mice were injected subcutaneously with analgesics (5 mg/kg ketoprofen, 0.1 mg/kg buprenorphine) and 1 mL of lactated Ringer’s. The mice were shaved in the area over and surrounding the spleen and the shaved area was disinfected with 70% ethanol and iodine. The abdominal cavity was opened, and the spleen was removed by cauterizing the splenic arteries. The abdominal cavity and skin were closed using 4-0 gauge synthetic absorbable suture. The mice were then treated with ketoprofen (s.i.d.) and buprenorphine (b.i.d.) for 3 days post-surgery and allowed to recover 4 weeks prior to the initiation of treatments and diet.

#### Preparation of EOCs and CM for in vivo studies

Treated EOCs were detached using Accutase Enzyme Detachment Buffer (eBioscience, San Diego, CA) washed once, and resuspended in saline for injections. CM was concentrated 10-fold using a < 3 kDa Amicon centrifuge filtration column (Millipore, Etobicoke, ON, Canada) and dialyzed twice in saline prior to injection.

#### Treatment and sacrifice of Ldlr-/- mice

Starting at 8 weeks of age, *Ldlr*-/- mice were fed a Western style diet (#88137 Harlan Teklad) for 8 weeks. The splenectomized cohort of the *Ldlr*-/- mice were injected via the tail vein with 10^6^ EOCs from WT C57Bl/6 mice in 100 μL of saline. The non-splenectomized *Ldlr*-/- mice received 100 μL of the 10-fold concentrated CM. Treatments were administered every 2 weeks throughout the Western diet feeding period. At sacrifice, the mice were exsanguinated under isoflurane anesthesia and were perfused with 10 mL of 2% EDTA/PBS. The heart was placed in 4% paraformaldehyde for 2 hours. The heart was transferred to 30% sucrose overnight at 4°C and then embedded in OCT (Cedarlane, Burlington, ON, Canada) for sectioning. Whole blood obtained at sacrifice was centrifuged at 500*xg* for 20 minutes at 4°C for the separation of plasma, which was aliquoted and frozen at -80°C until analysis. Plasma cholesterol was determined by enzymatic assay using the Cholesterol Infinity Kit (Thermo Scientific) as per the manufacturer’s recommendations.

#### Histological analysis

Aortic sinuses were cryosectioned using a Leica CM 3050S cryostat (Concord, ON, Canada) at a 10 μm thickness and placed on Superfrost Plus microscope slides (Fischer Scientific). The slides were stored at -80°C until use. Sections were dried at room temperature prior to staining. The sections were washed 3 times in water and then stained with hematoxylin (Sigma-Aldrich) for 5 minutes. After 3 water washes, the sections were dipped in acid alcohol for 10 seconds, followed by 3 water and 1 PBS wash, and then stained in 0.5% eosin solution (Sigma-Aldrich) for 3 minutes. After staining, the sections were dehydrated in sequential alcohol washes for 3 minutes each, followed by 3 washes in toluene for 1 minute each. The sections were then mounted for imaging. Images were acquired using a Mirax Scan slide scanner (Zeiss). Sections were analyzed by a scientist blinded to the treatment conditions. Lesion area is reported as a percent of total lumen area.

#### Statistical analysis

One-way ANOVA, followed by the Holm’s Sidak’s post hoc test, was used for comparison between more than two groups. For the human EOC study, no sample size calculation was performed as the analysis was considered exploratory. A parametric paired 2-tailed *t* test was used to compare treatment groups in the human EOC studies. A value of *P*<0.05 was considered to be statistically significant. Statistical tests were performed using GraphPad Prism (GraphPad Software Inc., La Jolla, CA).

## Supplemental Figures

**Supplemental Figure 1.**
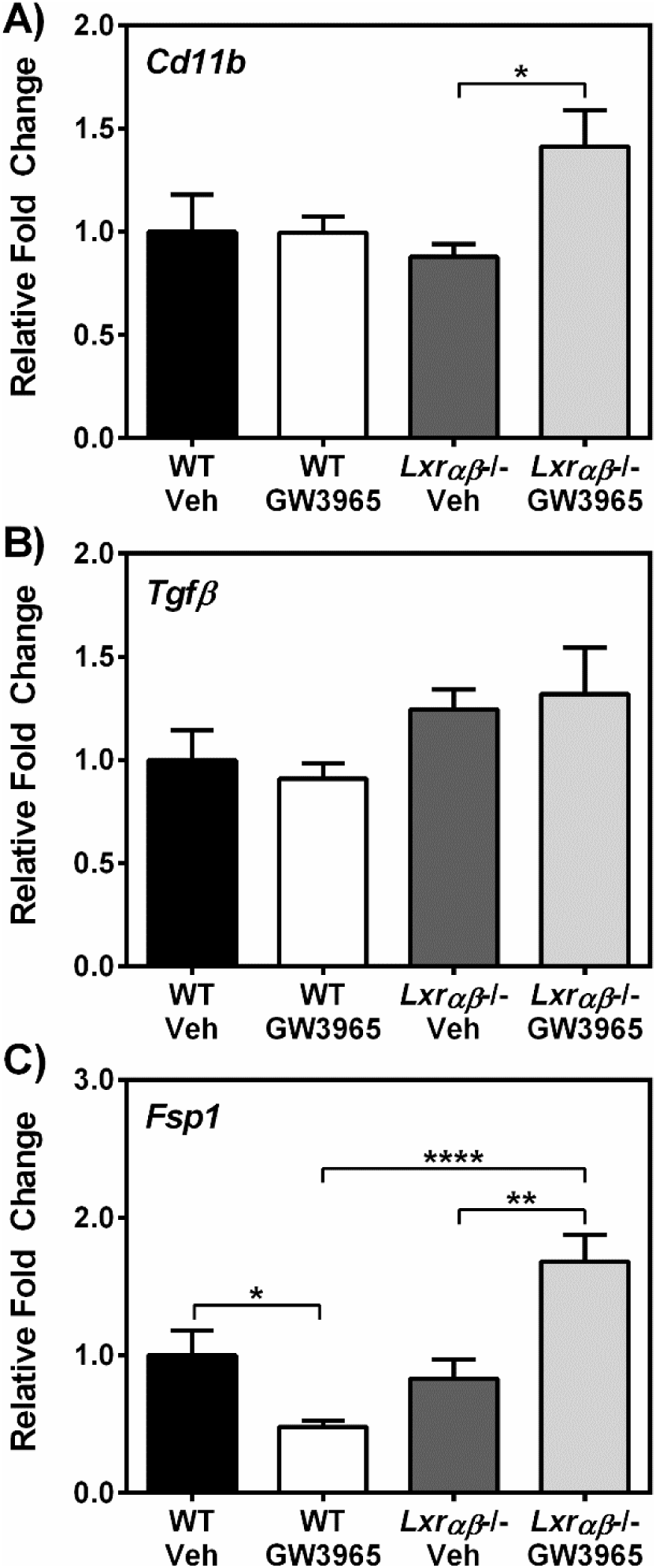
Differentiation of EOCs in the presence of GW3965 does not induce myeloid or mesenchymal markers. EOCs from WT and *Lxrαβ*-/- mice were treated with GW3965 and gene expression was performed for **A**, *Cd11b*, **B**, *Tgfβ*, and **C**, *Fsp1*. n=6 per group. Data represent the mean ± SEM. \**P*<0.05, \*\**P*<0.01, \*\*\**P*<0.001, \*\*\*\**P*<0.0001.

**Supplemental Figure 2.**
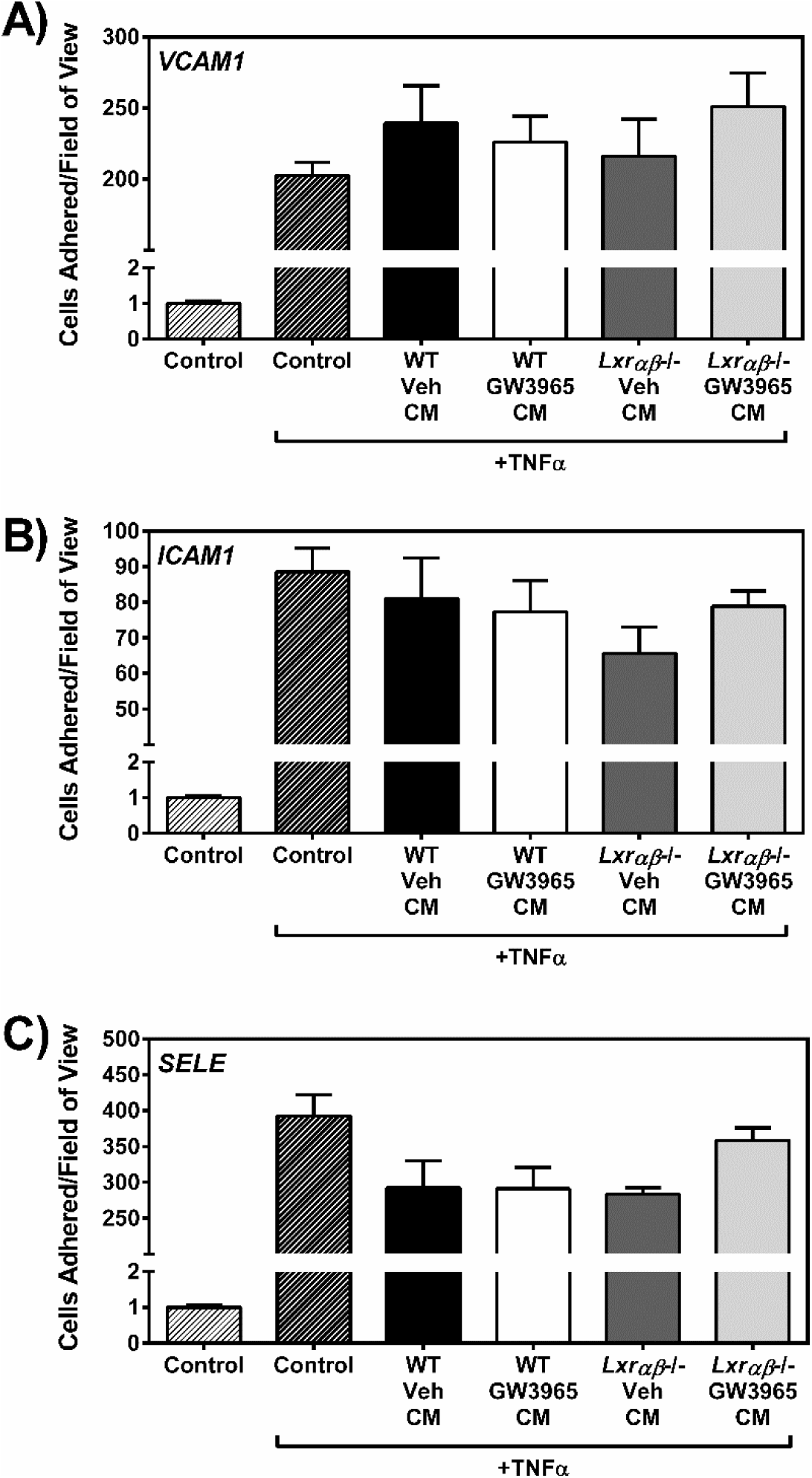
Incubation with the secretome from GW3965-treated EOCs does not reduce the expression of adhesion molecules and selectins on activated endothelial cells. HUVECs were incubated with conditioned media from treated EOCs and gene expression was assessed for **A**, *VCAM1*, **B**, *ICAM1*, and **C**, *SELE*. n=6 per group. Data represent the mean ± SEM.

## Supplemental Tables

**Supplemental Table 1.**
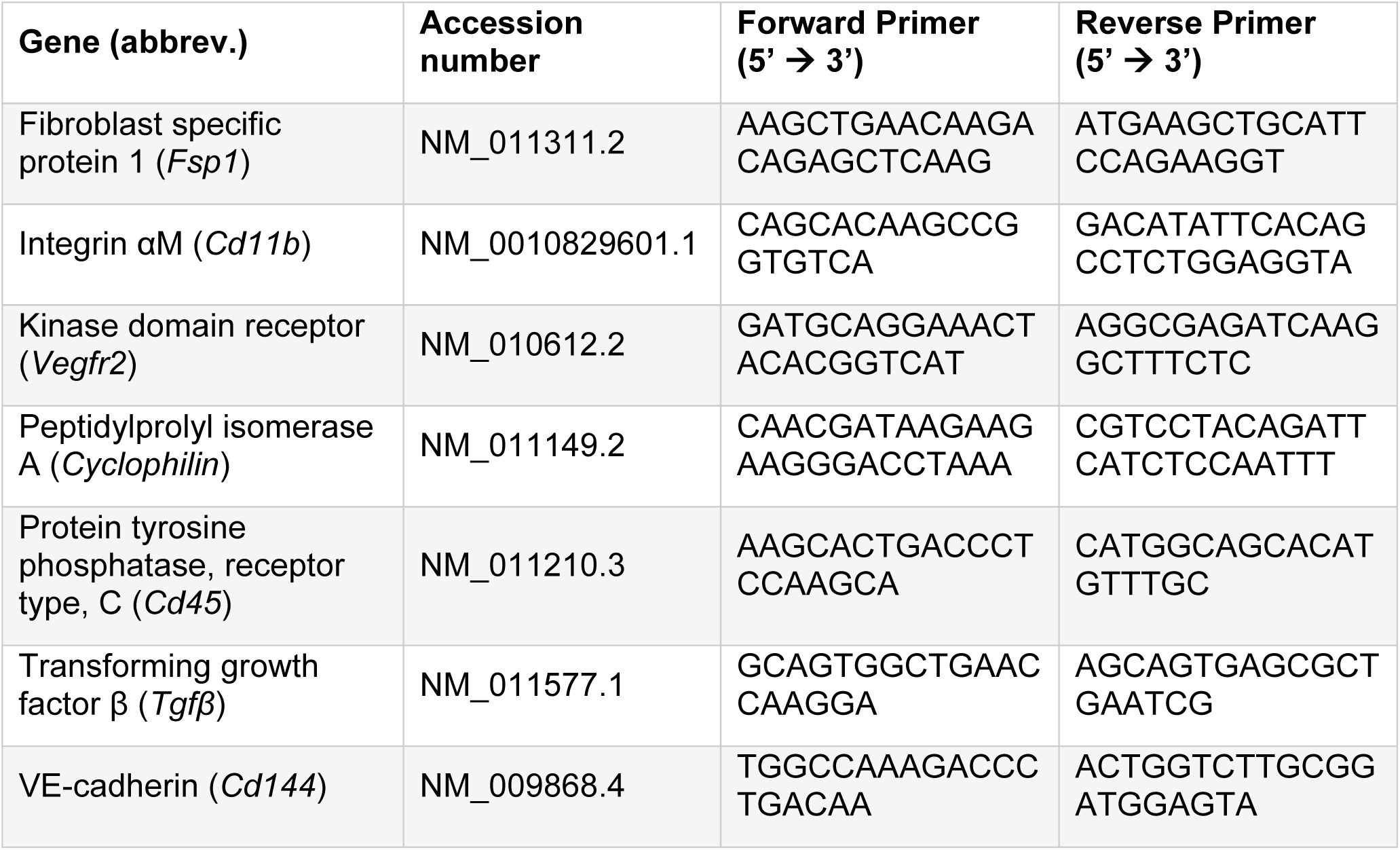
List of mouse primers.

**Supplemental Table 2.**
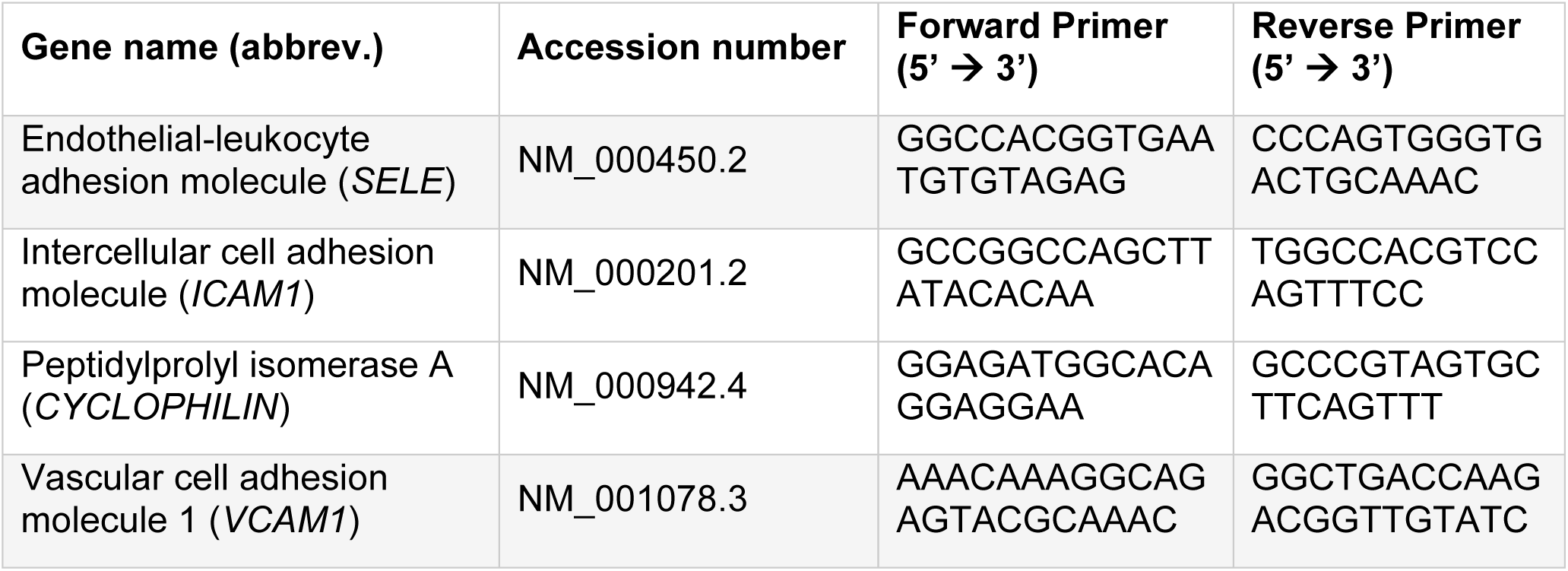
List of human primers.

**Supplemental Table 3.** Circulating immune cells in splenectomized *Ldlr*-knockout mice receiving EOCs.

|  | **Saline** | **Veh EOC** | **GW3965 EOC** |
| --- | --- | --- | --- |
| RBC ($10^{12}$ cells/L) | 10.0 ± 0.3 | 9.9 ± 0.2 | 10.4 ± 0.1 |
| WBC ($10^9$ cells/L) | 13 ± 1 | 12 ± 1 | 13.0 ± 0.8 |
| Lymphocyte ($10^9$ cells/L) | 10 ± 3 | 10 ± 1 | 10.1 ± 0.6 |
| Platelet ($10^9$ cells/L) | 590 ± 40 | 670 ± 20 | 610 ± 30 |
| Monocyte ($10^9$ cells/L) | 0.55 ± 0.06 | 0.59 ± 0.05 | 0.59 ± 0.05 |
| Neutrophil ($10^9$ cells/L) | 2.7 ± 0.4 | 2.1 ± 0.1 | 2.2 ± 0.4 |

n=10-11 mice per group. Data represent the mean ± SEM. RBC: red blood cell, WBC: white blood cell.

**Supplemental Table 4.**
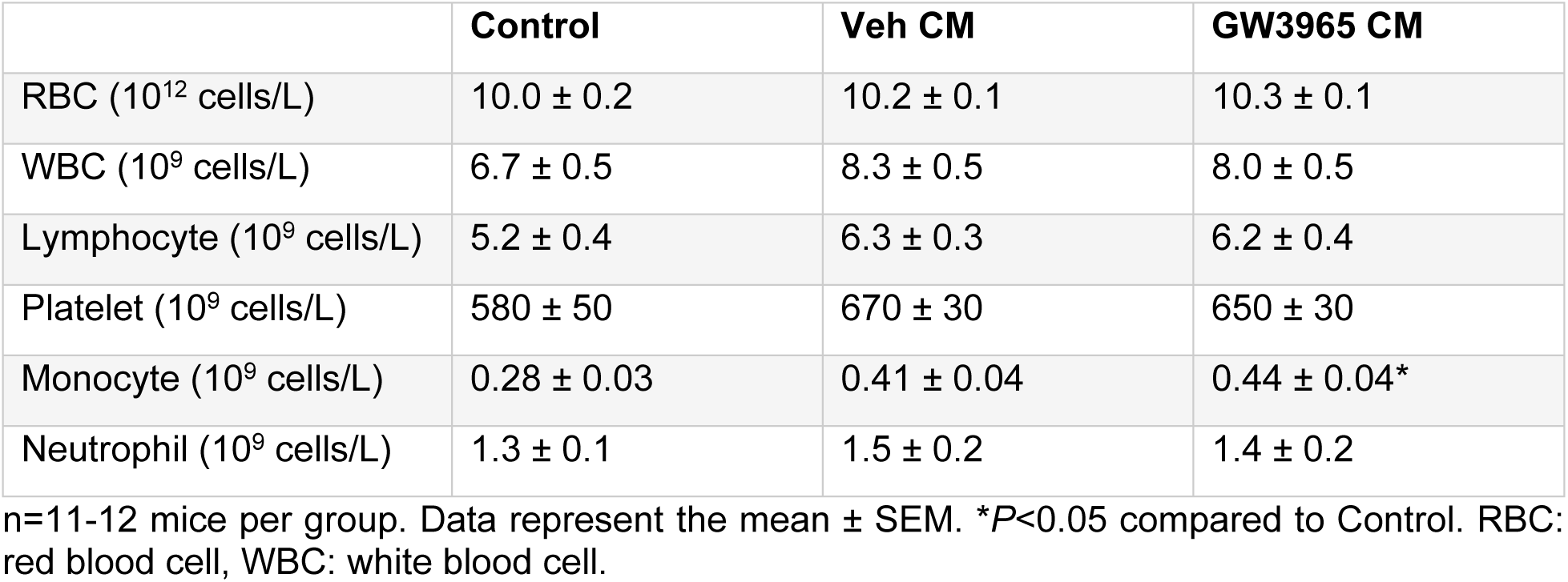
Circulating immune cells in *Ldlr*-knockout mice receiving conditioned media derived from treated EOCs.

|  | **Control** | **Veh CM** | **GW3965 CM** |
| --- | --- | --- | --- |
| RBC ($10^{12}$ cells/L) | 10.0 ± 0.2 | 10.2 ± 0.1 | 10.3 ± 0.1 |
| WBC ($10^9$ cells/L) | 6.7 ± 0.5 | 8.3 ± 0.5 | 8.0 ± 0.5 |
| Lymphocyte ($10^9$ cells/L) | 5.2 ± 0.4 | 6.3 ± 0.3 | 6.2 ± 0.4 |
| Platelet ($10^9$ cells/L) | 580 ± 50 | 670 ± 30 | 650 ± 30 |
| Monocyte ($10^9$ cells/L) | 0.28 ± 0.03 | 0.41 ± 0.04 | 0.44 ± 0.04* |
| Neutrophil ($10^9$ cells/L) | 1.3 ± 0.1 | 1.5 ± 0.2 | 1.4 ± 0.2 |

n=11-12 mice per group. Data represent the mean ± SEM. \* $P$ <0.05 compared to Control. RBC: red blood cell, WBC: white blood cell.

